# CB1 and CB2 receptors differentially modulate the cognitive impact of maternal immune activation and perinatal cannabinoid exposure

**DOI:** 10.1101/2024.11.16.623455

**Authors:** Han-Ting Chen, Ken Mackie

## Abstract

Maternal immune activation (MIA) commonly arises in response to an infection during pregnancy. MIA elevates cytokine levels, triggering an inflammatory cascade, which may be detrimental to the developing nervous system. Similarly, cannabis use and exposure of the fetus to cannabinoids during pregnancy (PCE) may elicit neuroinflammation and lead to detrimental behavioral outcomes. This is particularly concerning as there has been a notable rise in cannabis use during pregnancy. This study endeavors to examine the interaction between MIA and PCE and elucidate the role of CB1 and CB2 receptors in MIA and PCE outcomes. To this end, we compared the impact of MIA, PCE and MIA+PCE in wildtype, CB1, and CB2 cannabinoid receptor knockout mice of both sexes. PCE was modeled by daily 3 mg/kg THC administration from gestational day 5 (GD5) to postnatal day 10. MIA was modeled by intravenous Poly (I:C) injection at GD16.5. Subsequently, we assessed emotional and cognitive behaviors of adult offspring. Adult male offspring of dams exposed to PCE or MIA were impaired in novel object recognition and the delayed alternation working memory tasks. Interestingly, these behavioral impairments were absent when MIA and PCE were combined. Cannabinoid receptor knockout studies found that CB1 receptors mediated behavioral deficits after PCE. In contrast CB2 receptors were necessary for full expression of MIA-induced behavioral impairments. Although females showed more modest behavioral changes after MIA or PCE, CB1 receptors were required for the PCE deficit and CB2 receptors were required for the MIA deficit also in females. Notably, lack of CB2 receptors in males prevented the “protection” following combined MIA + PCE, while CB1 knockout mice remained protected. Taken together, these results suggest a complex interplay between PCE, MIA and CB1 and CB2 cannabinoid receptors.

**Highlights:**

- Both PCE and MIA impair cognitive behaviors.
- Combined PCE and MIA did not affect the cognitive behaviors examined.
- CB1 receptors are required for deficits after prenatal cannabis exposure.
- CB2 receptors are required for deficits after maternal immune activation.
- CB2 receptors are necessary for protection from deficits by combined PCE and MIA.

## 1. Introduction

Legalization of cannabis for non-medical use in the United States started in 2012 through voter initiatives in Colorado and Washington. Currently, cannabis is legalized in 14 states for medical use and in an additional 25 states for both medical and recreational use. In states with legalized cannabis, the percentage of young adults (19-30 years old) reporting past-year cannabis use was approximately 44% of those surveyed in 2022, increasing from five (35% in 2017) and 10 (28% in 2012) years ago (1). Daily cannabis use in this population also steadily increased, reaching its highest level in 2022 (11%), which was greater than five (8% in 2017) and 10 (6% in 2012) years ago (1). Thus, the incidence of cannabis use in the young adult population has been steadily increasing over the past 10 years. Focusing on cannabis use during pregnancy, the number of women reporting use during pregnancy doubled from 2002 to 2014, an increase that preceded widespread legalization (2). More recently during the COVID pandemic, cannabis use during pregnancy further increased by 25% over pre-pandemic levels, as assessed by an interrupted-time-series model (3). More details about patterns of cannabis use during the different stages of pregnancy have recently emerged. After legalization (2016–2017), overall cannabis uses in pregnant women increased from 3.4% to 7.0%. Examining cannabis use during specific trimesters, use ranged from 5.7% to 12.1% during the first trimester, from 0.6% to 2.5% during the second trimester and from 0.5% to 2.5% during the third trimester (4). The higher levels of cannabis use during the first trimester are likely due to use before the pregnancy is known to the mother and because cannabis products may be suggested as a treatment of nausea in the first trimester (e.g., as recommended by some cannabis dispensaries (5)).

Concerningly, there is a growing body of evidence that perinatal cannabinoid exposure (PCE), such as would occur from cannabis use during pregnancy, may affect mental health in the offspring (6). For example, a study followed maternal cannabis users and assessed their children’s behavior at 5 years old. These researchers observed that the cannabis-exposed children showed more aggressive behaviors, attention deficit/hyperactivity problems, and oppositional/defiant behaviors, as well as decreased cognitive flexibility and weaker receptive language, even after correcting for numerous potential confounders, including nicotine exposure (7). A preclinical study using rats found that male offspring of PCE dams had decreased prepulse inhibition (PPI), hyperdopaminergic function, and extensive alterations in dopamine synapses (8) and other studies have found general cognitive deficits in the offspring (9–12). Additional PCE effects include impaired response to fluoxetine in the forced swim test (13). Activating the endocannabinoid system during pregnancy also affects offspring behaviors. For example, perinatal exposure to URB597, which would increase the endocannabinoid, anandamide, increased depressive behaviors and impaired working memory (14). Moreover, both prenatal and adolescent cannabis can affect the immune system (15). Prenatal effects may occur as THC crosses the fetal–placental barrier to affect fetal neurons, glia, and immunocytes. Prenatal effects may also be mediated by maternal lymphocytic CB1 and CB2 receptors that are positioned to modulate the anti-inflammatory cytokine IL-10 (16). Thus, cannabinoids may exert their effects on the developing nervous system directly, via CNS cannabinoid receptors or indirectly via their effects on immune cells.

Viral infection and the subsequent maternal immune activation (MIA) are strongly associated with deficits in fetal neurodevelopment and may cause severe and long-lasting damage to the CNS (17–19). For example, individuals whose second trimester of fetal development corresponded to the 1957 influenza pandemic had a higher risk of being diagnosed with schizophrenia and being admitted to a psychiatric hospital (20). Several preclinical models of MIA have been developed. Typically, they avoid using infectious materials, instead relying on indirect activation of the innate immune system. Different activators of the innate immune system during pregnancy also lead to different consequences in these model systems. For example, treating mice with Lipopolysaccharides (LPS) on the 14^th^ day of pregnancy, to mimic gram-negative bacterial infection, increased levels of maternal inflammatory cytokines, including IL-17a, and corticosterone. LPS treatment of the dam also increased anxiety behaviors in juvenile offspring and impaired several measures of social behavior in adults (21). Poly(I:C) is often used to model viral infection in preclinical studies, activating innate immune pathways by mimicking viral nucleic acids (22). Administering poly(I:C) to mice on gestational day 15^th^ increases serum IL-6 levels within 3 hours and increases microglia activation in P21 male offspring (23). In addition to the biochemical changes, prenatal poly(I:C) also decreased PPI in adult offspring (18,24), which was accompanied by microglia activation in the hippocampus and corpus callosum (25). Giving poly(I:C) during late gestation also reduced working, cross-modal and visual memory and decreased sociability (25–27) in adult male offspring.

As reviewed above, both prenatal cannabis exposure and viral infection may adversely affect the developing CNS. Given the widespread and increasing use of cannabis during pregnancy and the high prevalence of viral infections during pregnancy, an important question is: “Does a viral infection in the setting of prenatal cannabis use more severely impair CNS development?” To address this question, we designed an experimental paradigm based on a neurodevelopmental time line (17) that mimics an individual chronically using cannabis during pregnancy who subsequently experiences a viral infection, to see whether this would worsen the behavioral performance of the offspring. Unexpectedly, we found that although, individually, MIA (elicited by poly(I:C)) and PCE (mimicked by THC injection) both impaired several domains of offspring cognitive behavior, combining treatments prevents this impairment. Further experiments in mice lacking CB1 or CB2 receptors suggest the protective and detrimental effects of THC, respectively, are mediated by different cannabinoid receptors, possibly through their differential regulation of neuroinflammatory responses.

## 2. Material and Methods

### 2.1. Husbandry

Wildtype, CD1 mice (IGS) were purchased from Charles River (Strain 022). The CB1 line was provided by Catherine Ledent (28) and has been maintained on a CD1 background in our laboratory for almost 25 years. The CB2 line was purchased from JAX (Strain 005786, (29)), backcrossed onto a CD1 background. Both transgenic lines were annually backcrossed onto CD1s. To prepare treated dams in all 3 lines of mice used, trio-breeding cages were set up with one male and two 12-week-old females of the indicated genotypes. Once a vaginal plug was observed, the dam was moved to its own cage. The day the vaginal plug was found was denoted as gestational day (GD) 0.5. On postnatal day 21 (P21) pups, were weaned and separated based on sex with 2-4 housed per cage under constant temperature with a 12-hr light-dark cycle (07:00-19:00). Litter sizes and sex ratios were not affected by the treatments. Food and water were available ad *lib* except as specified in the experimental design. All experimental protocols were approved by the Indiana University Bloomington Institutional Animal Care and Use Committee.

### 2.2. Drug preparation and administration

THC (provided by NIDA Drug Supply) was diluted in 100% EtOH to a 6mg/ml stock solution. This stock was then mixed in a ratio of 1:1:18 with Cremophor EL and sterile normal saline and injected at a dose of 3 mg/kg subcutaneously from GD5.5 to P10 (PCE). Poly(I:C) (Sigma, P9586) was dissolved in sterile normal saline to 10mg/ml and on GD16.5 injected intravenously at 0.5ml/kg to give a dose of 5mg/kg. Vehicle administration for the THC and poly(I:C) groups was the weight-based volume of 1:1:18 solution or sterile saline, respectively. Pregnant dams were assigned into four different treatment groups: (1) a control group, receiving chronic vehicle and GD16.5 normal saline, (2) a MIA group, receiving chronic vehicle and poly (I:C) 5mg/kg on GD16.5, (3) a PCE group, receiving chronic THC 3mg/kg from GD5.5 to P10 and normal saline on GD16.5, modeling PCE with no viral infection, and (4) a MIA+PCE combined group, with chronic THC 3 mg/kg and poly(I:C) 5mg/kg given on the days described above. Pups received no treatment and were weaned at P21. Behavioral testing of offspring started at P50 to identify the effects of the various maternal treatments on offspring behavior.

### 2.3. Behavioral methods

Treatments were conducted on at least 2–3 dams per treatment condition across multiple litters. Results from the first two experimental cohorts were initially evaluated. If the performance of animals in the first two cohorts was inconsistent, a third cohort was added. As a result, we used two cohorts for wildtype and CB1KO mice, and three cohorts for CB2KO mice.

Behavioral experiments were initiated 2-3 hours into the light phase to avoid the hyperactivity exhibited during the dark/light transition and ended around 18:30 (dark cycle started at 19:00). Experiments were conducted under reduced illumination (100 lux) and all equipment was cleaned with disinfectant solution (Rescue 1:64, Oakville, ON, Canada, product# 23305) and 75% ethanol between tests to reduce any residual odors that might affect the following experimental subject.

When identifying cannabinoid receptor involvement in the behaviors, we compared CB1 and CB2 KO vehicle control mice with simultaneously treated wildtype control mice to ensure the experimental effectiveness of MIA and PCE in case the absence of cannabinoid receptors affected memory task performance. For example, knocking out CB1 receptors can enhance object recognition (30) and spatial memory may be affected by deletion of CB1 receptors from the hippocampus (31). Absence of CB2 receptors has been reported to enhance spatial memory, but impair long-term (32) and social (33) memory.

#### 2.3.1. Open field activity

The mouse was allowed to freely explore a plexiglass chamber (42L×42W×32H (cm)) for 15 minutes. The mouse’s movements, position in the chamber, and distance traveled were recorded and analyzed by EthovisionXT.

#### 2.3.2. Elevated plus maze

A 50-cm elevated plus maze with 5x30-cm arms, two open, two closed, 50 cm above the floor was used. The mouse was placed in the center-open area facing in a random direction. The mouse’s movements were recorded for 6 minutes. Speed, entries into each arm and amount of time spent in each arm and the center area were recorded and analyzed by EthovisionXT.

#### 2.3.3. Novel object recognition

The mouse to be tested was habituated to the testing arena by being placed in an empty plexiglass chamber (42L×42W×32H (cm)) for 5 minutes on the day of the test. To start the training phase, after habituation the mouse was returned to its home cage for 3 minutes and the chamber cleaned, and two identical objects placed into the chamber. The mouse was returned to the chamber and videotaped while freely exploring the chamber and objects for 10 minutes. The mouse was then returned to its cage for 2 hours. To test the mouse’s recognition memory, one familiar and one novel (different color and shape but similar in size and height) object were placed in a clean chamber. The mouse then explored the chamber and objects for 10 minutes. The mouse’s movements in the training and testing phases were recorded and analyzed by EthovisionXT to determine the attention given to the different objects. The DI value was defined as the difference in exploration time between novel and familiar objects, divided by the total exploration time for both objects. DI difference was the difference in individual DI values between the training (accounting for object preference shown during training) and testing phases.

#### 2.3.4. Working memory

A T-maze (6x30 cm) was made from plexiglass with gates at the intersection of each arm that could be closed to force the mouse to turn towards a specific arm. The experimental protocol was modified from (34,35). Animals were subjected to a daily 4-hour food restriction for a week before the training phase, and then had a 2-hour food restriction prior to testing for the remainder of the experiment. For the first two days of habituation, animals were able to freely explore the maze for 5 minutes, accompanied by their cage mates (day 1) or alone (day 2). For the next two days, forced-learning training was conducted with the gate on one side closed so the mouse could only enter the assigned arm to retrieve a food reward (cereal pellet). Delayed alternating response training started on the fifth day.

For the training, 10 trials were conducted daily, with each trial including a forced-choice and a free-choice. The forced-choice trial was conducted identically as the forced-learning training trial (above). After the forced-choice trial, the mouse was trapped in the start arm for 10 seconds before participating in the free-choice trial. The free-choice trial started with both doors open and the mouse was free to choose either arm, but only the arm opposite from that available during the forced-choice trial contained a reward (cereal pellet). After a 1-minute delay, the combination of forced and free trials was repeated. The daily success rate for each mouse was calculated as the percent of the ten trials that were correctly completed to assess their learning ability during the 8 days of training.

Memory retention was tested on the day following the last training session. The protocol of increased-delay testing was the same as the training protocol, except only five trials were conducted and we varied the internal delay time during which the mouse was trapped in the start arm between the forced and free choice trials from 10 seconds to 15, 30 and 60 seconds. Each trial was repeated in blocks of five and the success rate determined for each delay interval.

### 2.4. Statistical analysis

Behavioral data were analyzed using GraphPad Prism9 and are expressed as mean ± SEM. Normality was tested using the Shapiro-Wilk normality test. If the distribution of residuals was normal, two-way ANOVA was used to analyze overall significance for different behavioral experiments, with Sidak’s post hoc comparisons to determine significance. If the distribution of residuals was not normal, the Kruskal-Wallis test was used to determine significance. A p value less than 0.05 was accepted as statistically significant.

## 3. Results

### 3.1. PCE and MIA did not affect open field activity or anxiety behaviors

All mice were tested for their basic activity and anxiety levels prior to the cognitive behavioral tests. There were no significant differences in activity or anxiety levels between the different perinatal treatments or transgenic mouse strains (Table 1), indicating that the mice had similar levels of activity and anxiety prior to performing the cognitive behavioral tests.

**Table 1.**
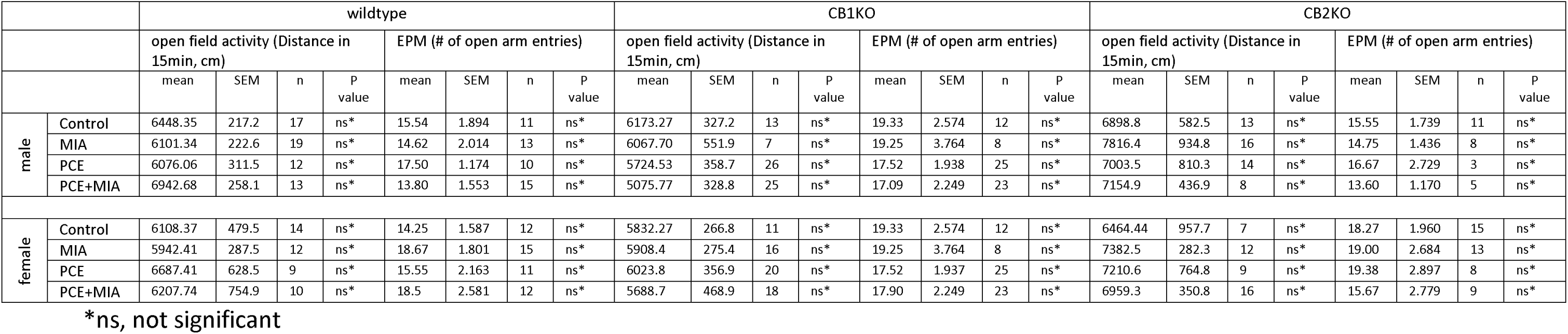
Impacts of MIA, PCE, and their combination on open field and elevated plus maze behaviors.

|  |  | wildtype |  |  |  |  |  |  |  | CB1KO |  |  |  |  |  |  |  | CB2KO |  |  |  |  |  |  |  |
| --- | --- | --- | --- | --- | --- | --- | --- | --- | --- | --- | --- | --- | --- | --- | --- | --- | --- | --- | --- | --- | --- | --- | --- | --- | --- |
|  |  | open field activity (Distance in 15min, cm) |  |  |  | EPM (# of open arm entries) |  |  |  | open field activity (Distance in 15min, cm) |  |  |  | EPM (# of open arm entries) |  |  |  | open field activity (Distance in 15min, cm) |  |  |  | EPM (# of open arm entries) |  |  |  |
|  |  | mean | SEM | n | P value | mean | SEM | n | P value | mean | SEM | n | P value | mean | SEM | n | P value | mean | SEM | n | P value | mean | SEM | n | P value |
| male | Control | 6448.35 | 217.2 | 17 | ns* | 15.54 | 1.894 | 11 | ns* | 6173.27 | 327.2 | 13 | ns* | 19.33 | 2.574 | 12 | ns* | 6898.8 | 582.5 | 13 | ns* | 15.55 | 1.739 | 11 | ns* |
|  | MIA | 6101.34 | 222.6 | 19 | ns* | 14.62 | 2.014 | 13 | ns* | 6067.70 | 551.9 | 7 | ns* | 19.25 | 3.764 | 8 | ns* | 7816.4 | 934.8 | 16 | ns* | 14.75 | 1.436 | 8 | ns* |
|  | PCE | 6076.06 | 311.5 | 12 | ns* | 17.50 | 1.174 | 10 | ns* | 5724.53 | 358.7 | 26 | ns* | 17.52 | 1.938 | 25 | ns* | 7003.5 | 810.3 | 14 | ns* | 16.67 | 2.729 | 3 | ns* |
|  | PCE+MIA | 6942.68 | 258.1 | 13 | ns* | 13.80 | 1.553 | 15 | ns* | 5075.77 | 328.8 | 25 | ns* | 17.09 | 2.249 | 23 | ns* | 7154.9 | 436.9 | 8 | ns* | 13.60 | 1.170 | 5 | ns* |
| female | Control | 6108.37 | 479.5 | 14 | ns* | 14.25 | 1.587 | 12 | ns* | 5832.27 | 266.8 | 11 | ns* | 19.33 | 2.574 | 12 | ns* | 6464.44 | 957.7 | 7 | ns* | 18.27 | 1.960 | 15 | ns* |
|  | MIA | 5942.41 | 287.5 | 12 | ns* | 18.67 | 1.801 | 15 | ns* | 5908.4 | 275.4 | 16 | ns* | 19.25 | 3.764 | 8 | ns* | 7382.5 | 282.3 | 12 | ns* | 19.00 | 2.684 | 13 | ns* |
|  | PCE | 6687.41 | 628.5 | 9 | ns* | 15.55 | 2.163 | 11 | ns* | 6023.8 | 356.9 | 20 | ns* | 17.52 | 1.937 | 25 | ns* | 7210.6 | 764.8 | 9 | ns* | 19.38 | 2.897 | 8 | ns* |
|  | PCE+MIA | 6207.74 | 754.9 | 10 | ns* | 18.5 | 2.581 | 12 | ns* | 5688.7 | 468.9 | 18 | ns* | 17.90 | 2.249 | 23 | ns* | 6959.3 | 350.8 | 16 | ns* | 15.67 | 2.779 | 9 | ns* |

\*ns, not significant

### 3.2. Effects of PCE and MIA on novel objection recognition

We first examined the effects of perinatal cannabinoid exposure (PCE) with or without maternal immune activation (MIA) on novel object recognition (NOR) in adult male offspring. Wildtype male mice showed significant novel objective recognition (Figure 1A, F (1, 24) = 13.36, P=0.0013). Sidak’s post hoc comparisons found that male offspring from vehicle-treated (perinatal vehicle treatment plus normal saline on E16.5) significantly recognized a novel object (P=0.0077, 95% C.I. = [-0.4655, -0.05291]). However, after MIA they failed to distinguish between the novel and familiar objects (P=0.9995, 95% C.I. = [-0.1913, 0.2213]). Similarly, PCE mice failed to distinguish between the two objects (P=0.3743, 95% C.I. = [-0.3371, 0.07550]).

**Figure 1.**
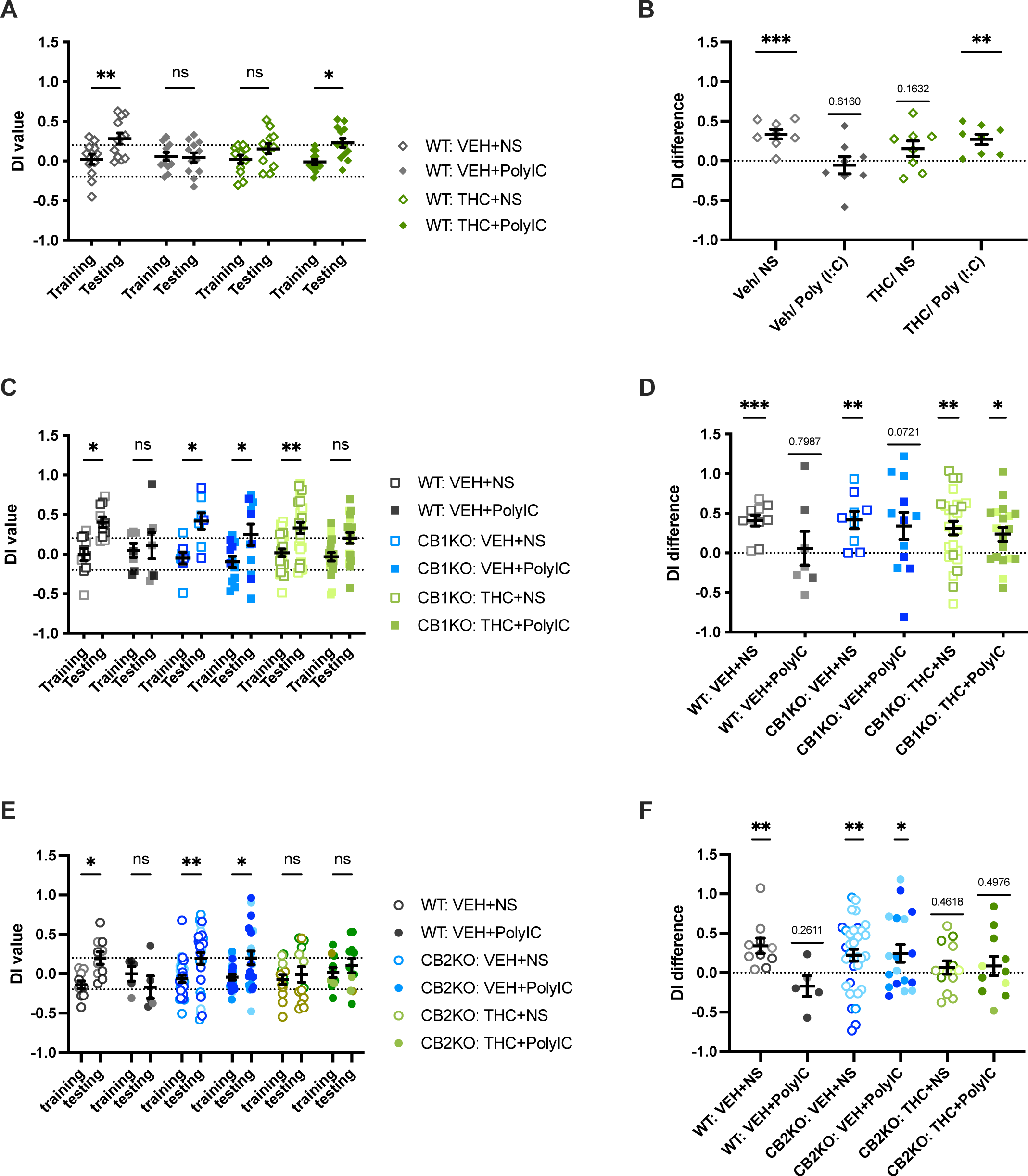
Maternal THC and MIA, but not their combination, impairs novel object recognition in male offspring. Prenatal treatments, novel objection recognition behavior and DI differences were performed and calculated as described in Methods for wildtype, CB1 KO, and CB2 KO mice. Wildtype mice (DI values in **A**, DI differences in **B**). CB1KO mice (DI values in **C**, DI differences in **D**). CB2KO mice (DI values in **E**, DI differences in **F**). Gray shad as wildtype, blue as male mice perinatal chronically treated vehicle and green as perinatal chronically treated THC. Opened symbols represent administration of normal saline to the dam at E16.5, while solid symbols represent administration of poly (I:C) at E16.5. CB1KO is square and CB2KO is circle. Sample size in panel A and B: 12 for wildtype vehicle/ns, 12 for wildtype vehicle/poly(I:C), 12 for wildtype THC/ns, 13 for wildtype THC/poly(I:C) ;in panel C and D: 10 for wildtype vehicle/ns, 7 for wildtype vehicle/poly(I:C), 8 for CB1KO vehicle/ns, 12 for CB1KO vehicle/poly(I:C), 26 for CB1KO THC/ns, 19 for CB1KO THC/poly(I:C); in panel E and F: 10 for wildtype vehicle/ns, 5 for wildtype vehicle/poly(I:C), 29 for CB2KO vehicle/ns, 21 for CB2KO vehicle/poly(I:C), 13 for CB2KO THC/ns, 11 for CB2KO THC/poly(I:C). Levels of significance: *, p<0.05; **, p<0.01; ***, p<0.001 compared with chronic vehicle and normal saline (A, C, and E) or compared to a DI difference of 0 (B, D, and F).

Surprisingly, mice retained their ability to discriminate between the two objects if PCE and MIA treatments were combined (P=0.0110, 95% C.I. = [-0.4382, -0.04181]). To better address the differences in individual mice, we investigated each mouse’s DI value difference (i.e., the change in object discrimination between the two familiar objects and the novel/familiar object for each mouse). Control male offspring showed a significant ability to distinguish the two objects after training (Figure 1B, t (7) =5.575, P=0.0008), while the MIA (t (7) =0.5246, P=0.6160) and PCE offspring (t (7) =1.558, P=0.1632) did not. As with the group DI differences, combining PCE and MIA resulted in retention of object discrimination (t (7) =4.164, P=0.0042). Surprisingly, in female offspring, the wildtype vehicle control mice couldn’t reliably discriminate between the novel and familiar objects under our experimental conditions (Figure 2A, F [3, 43] =14.38, P=0.2448), so if MIA and PCE affected this ability in this group of wildtype female offspring could not be determined.

**Figure 2.**
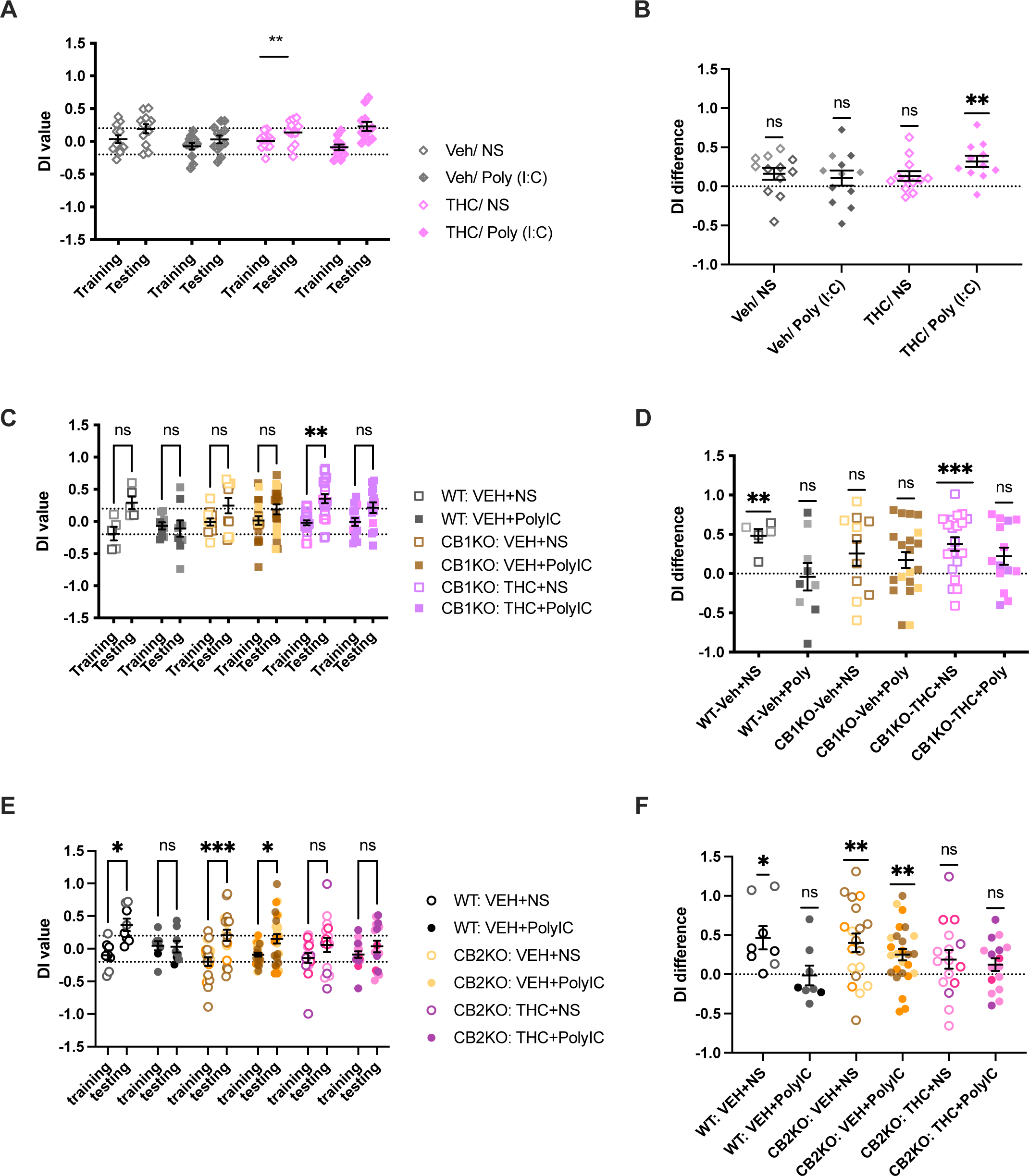
Maternal THC and MIA, but not their combination, impairs novel object recognition in female offspring. Prenatal treatments, novel objection recognition behavior and DI differences were performed and calculated as described in Methods for wildtype, CB1 KO, and CB2 KO mice. Wildtype mice (DI values in **A**, DI differences in **B**). CB1KO mice (DI values in **C**, DI differences in **D**). CB2KO mice (DI values in **E**, DI differences in **F**). Gray shad as wildtype, yellow as female as mice perinatal chronically treated vehicle, and purple as female offspring perinatal chronically treated THC. Opened symbols represent administration of normal saline to the dam at E16.5, while solid symbols represent administration of poly (I:C) at E16.5. CB1KO is square and CB2KO is circle. Sample size in panel A and B: 12 for wildtype vehicle/ns, 12 for wildtype vehicle/poly(I:C), 12 for wildtype THC/ns, 11 for wildtype THC/poly(I:C) ;in panel C and D: 5 for wildtype vehicle/ns, 9 for wildtype vehicle/poly(I:C), 11 for CB1KO vehicle/ns, 20 for CB1KO vehicle/poly(I:C), 20 for CB1KO THC/ns, 14 for CB1KO THC/poly(I:C); in panel E and F: 8 for wildtype vehicle/ns, 8 for wildtype vehicle/poly(I:C), 16 for CB2KO vehicle/ns, 26 for CB2KO vehicle/poly(I:C), 16 for CB2KO THC/ns, 16 for CB2KO THC/poly(I:C). Levels of significance: *, p<0.05; **, p<0.01; ***, p<0.001 compared with chronic vehicle and normal saline (A, C, and E) or compared to a DI difference of 0 (B, D, and F).

Next, we used knockout mice to determine the contributions of the two major cannabinoid receptors (CB1 and CB2) to the effects of PCE and MIA on object discrimination in male mice. Wildtype male mice receiving vehicle PCE were able to distinguish between the two objects (P=0.0233, 95% C.I. = [-0.7805, -0.03775]), as expected. As before, MIA in wildtype mouse impaired recognition ability (P=0.9996, 95% C.I. = [-0.5015, 0.3862]). Vehicle-treated CB1 knock out male mice were able to distinguish the novel from the familiar object (Figure 1C, F (1, 76) = 32.51, P<0.0001) similar as wildtype vehicle control (P=0.0193, 95% C.I. = [-0.8824, 0.05204]). The absence of CB1 receptors protected against loss of recognition memory after both MIA (P=0.0474, 95% C.I. = [-0.6805, -0.002506]) and PCE (P=0.0026, 95% C.I. = [-0.5438, -0.08316]) (Figure 1C, D). However, when MIA and PCE were combined in CB1 KO mice, the ability to distinguish between the two objects was lost when measured by group DI values (P=0.1156, 95% C.I. = -0.5059, 0.03294]) but persisted when measured by individual DI differences (Figure 1D; t (18) =2.724, P=0.0139). This suggests CB1 receptors are involved the adverse effects of MIA and PCE in recognition memory in male mice.

In female mice by 3 way ANOVA there was a significant effect by trial (Figure 2C, F [1, 61] =21.14, P<0.0001), but not by treatment (F [3, 61] =0.4712, P=0.7034). After multiple comparisons, the female CB1KO PCE group performed better (P=0.0011, 95% C.I. = [-0.6263, - 0.1268]) than did the CB1KO vehicle group by group DI values. In addition, analyzed individually the CB1KO PCE-treated female mice also showed significant recognition ability by DI differences (Figure 2D, t (19) =4.316, P=0.0004). These results suggest that CB1 receptors are involved in the effect of PCE on NOR performance in female mice. However, there were no differences in MIA and combined treated group by group DI values (Figure 2C, MIA, P=0.2859, 95% C.I. = [-0.4226, 0.07694], PCE+MIA, P=0.2292, 95% C.I. = [-0.5188, 0.07825]) and individual DI differences (Figure 2D, MIA, t (19) =1.718, P=0.1020, PCE+MIA, t (13) =2.020, P=0.0645) in female CB1 KO mice.

We next investigated the role of CB2 receptors in NOR impairment after PCE and MIA using CB2 receptor knock out mice. In these experiments, new sets of wildtype control mice were evaluated at the same time to validate the effectiveness and consistency of the experimental manipulations. Male CB2 KO mice were able to discriminate between the novel and familiar objects (Figure 1E, P=0.005, 95% C.I. = [-0.4563, -0.05798]; Figure 1F: DI difference, t (32) =2.883, P=0.0070) as did simultaneously evaluated WT mice (P=0.049, 95% C.I. = [-0.6796, -0.001334]; DI difference, t (9) =3.584, P=0.0059). However, while the WT male MIA offspring lost this ability (P=0.917, 95% C.I. = [-0.3089, 0.6504]; DI difference, t (4) =1.308, P=0.2611), CB2 KO male MIA offspring were able to distinguish between the two objects (P=0.037, 95% C.I. = [-0.4775, -0.009402]; DI difference, t (9) =3.584, P=0.0059). Both male CB2 KO (P=0.993, 95% C.I. = [-0.3622, 0.2328]; DI difference, t (12) =0.7603, P=0.4618) PCE offspring and CB2 KO mice receiving PCE + MIA failed to distinguish between the novel and familiar object (P=0.980, 95% C.I. = [-0.4085, 0.2382]; DI difference, t (10) =0.7038, P=0.4976). These results suggest that in male mice CB2 receptors are involved in the detrimental effects of MIA on recognition memory and are also involved in the protection on recognition offered by combined PCE + MIA. For female mice, new sets of wildtype control mice were tested. These mice discriminated between familiar and novel objects (Figure 2E, F [1, 87] = 24.83, P<0.001). CB2KO vehicle and MIA mice also showed significant recognition ability (control, P<0.001, 95% C.I.= [-0.6591, -0.1447]; MIA, P=0.019, 95% C.I.= [-0.4586, -0.02705]), while PCE wildtype and CB2 KO mice still showed impairment (P=0.330, 95% C.I.= [-0.4908, -0.08812]). As with CB2KO male mice, combined treatment impaired object recognition (Figure 2E, P=0.804, 95% C.I.= [-0.4041, 0.1565]). Similar results were found with individual DI differences: CB2KO control and MIA recognized the novel object (Figure 2F, vehicle control, t (18) =3.435, P=0.0030; MIA, t (25) =3.457, P=0.0020), and this ability was lost in the PCE and combined groups (PCE, t (15) =1.611, P=0.1280; PCE+MIA, t (15) =1.557, P=0.1403).

### 3.3. Effects of PCE and MIA on establishment of working memory

Working memory is an important form of memory to accomplish many daily tasks. To determine the impact of PCE and MIA on working memory we measured the rate of learning in the alternating T maze. Wildtype, CB1 KO, and CB2 KO mice were evaluated.

All four treatment groups of male wild type mice improved their performance with training (Figure 3A, F (7, 203) = 17.32, P<0.0001). However, the degree of improvement significantly varied by treatment (F (3, 29) =8.419, P=0.0004). Comparing the performance between groups, the MIA group started showing significantly impairment from vehicle control (F [1, 14] =19.06, P=0,0006) from day 3 (P=0.0273, 95% C.I. = [0.1879, 4.062]) and day 5-7 (day5, P=0.0485, 95% C.I. = [0.008717, 3.491]; day 6, P=0.0078, 95% C.I. = [0.4041, 3.096]; day 7, P=0.0078, 95% C.I. = [0.6259, 4.374]). Similarly, PCE mice performed worse than vehicle control mice (F [1, 14] =16.39, P=0.0012). However, like the NOR task, combining MIA with PCE significantly improved performance during training (F [4.491, 67.37] =15.31, P<0.0001) with no significant differences in learning between the MIA+PCE and vehicle control groups (F [1, 15] =1.579, P=0.2281). In wildtype female mice, all groups of mice learned over the training period (Figure 4A, F [7, 189] = 7.949, P<0.0001) and treatments showed a significant difference by 2-way ANOVA analysis (F [3, 27] = 13.64, P<0.0001). Compared to the control group, the MIA group was significantly impaired (F [1, 14] = 31.86, P<0.0001), as was the PCE group (F [1, 14] = 5.814, P=0.0302). However, performance after combining PCE + MIA was indistinguishable from the control group (F [1, 14] = 8.516e-005, P=0.9928), indicating female and male wildtype mice were similarly impacted by the various treatment conditions.

**Figure 3.**
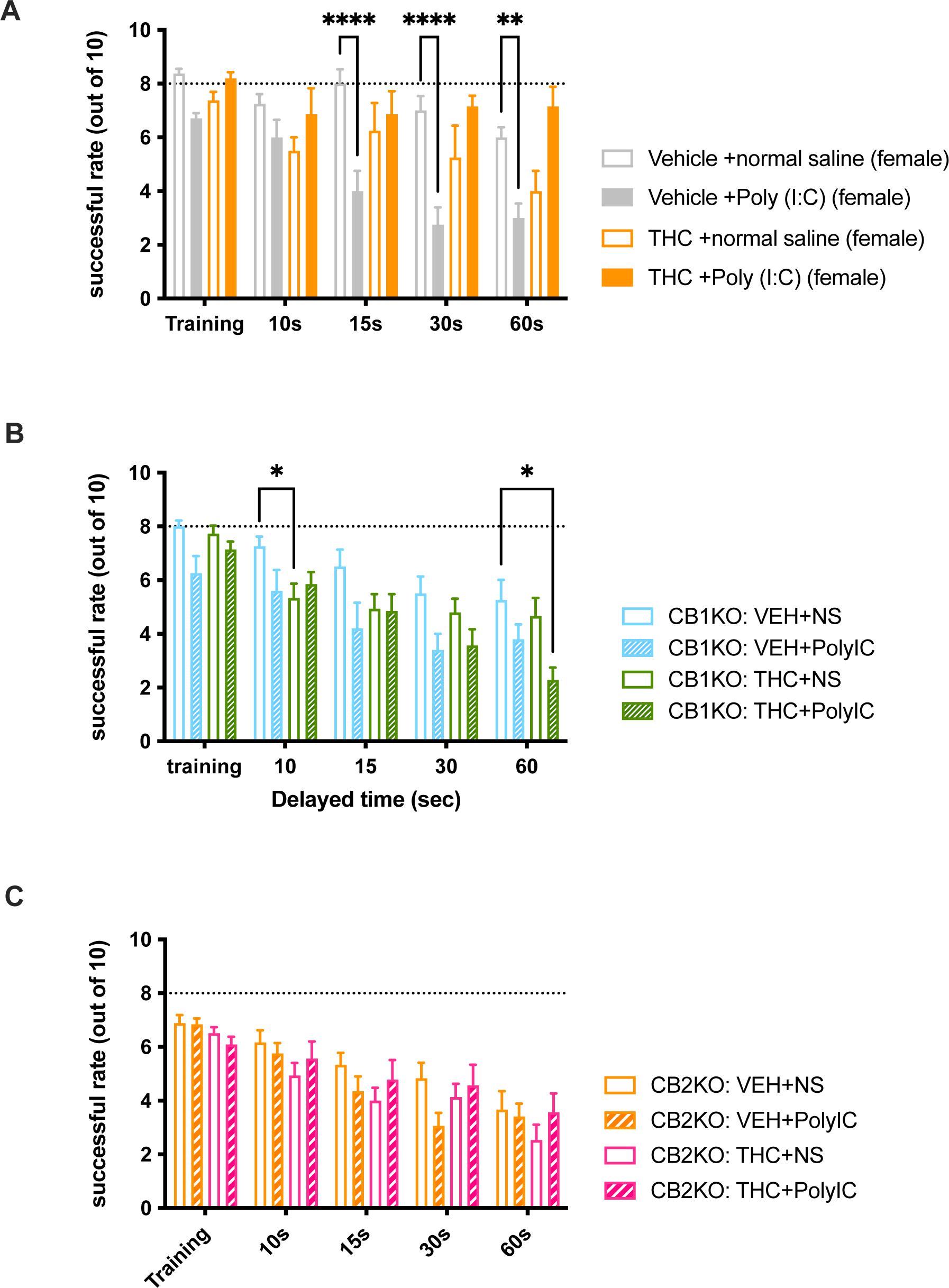
Maternal THC or MIA, but not their combination, slows learning in a working memory task in male offspring. **A.** Male offspring of wildtype dams treated with either THC or MIA learned the T maze task more slowly, but offspring of mice receiving THC and subjected to MIA learned the task as quickly as vehicle-treated mice. **B.** Maternal THC no longer impaired T maze learning in male CB1 KO offspring. However, MIA still impaired learning in CB1 KO offspring and combining THC and MIA rescued impairment. **C.** Maternal THC and MIA impaired T maze performance and THC no longer rescued performance in CB2 knockout mice. T maze learning was assessed by 8-days of daily training with fixed delayed and success rate recorded as a learning curve. Open symbols represent male offspring from dams receiving normal saline and solid symbols represent male offspring from dams receiving poly (I:C), both at GD16.5. Sample size in panel A: 8 for wildtype vehicle/ns, 8 for wildtype vehicle/poly(I:C), 8 for wildtype THC/ns, 9 for wildtype THC/poly(I:C) ;in panel B: 5 for wildtype vehicle/ns, 4 for wildtype vehicle/poly(I:C), 10 for CB1KO vehicle/ns, 8 for CB1KO vehicle/poly(I:C), 13 for CB1KO THC/ns, 14 for CB1KO THC/poly(I:C); in panel C: 12 for wildtype vehicle/ns, 5 for wildtype vehicle/poly(I:C), 13 for CB2KO vehicle/ns, 14 for CB2KO vehicle/poly(I:C), 9 for CB2KO THC/ns, 11 for CB2KO THC/poly(I:C)). *, p<0.05; ***, p<0.001 compared with chronic vehicle and GD16.5 normal saline; #, p<0.05; ###, p<0.001 compared with chronic vehicle and Poly (I:C) at GD16.5.

**Figure 4.**
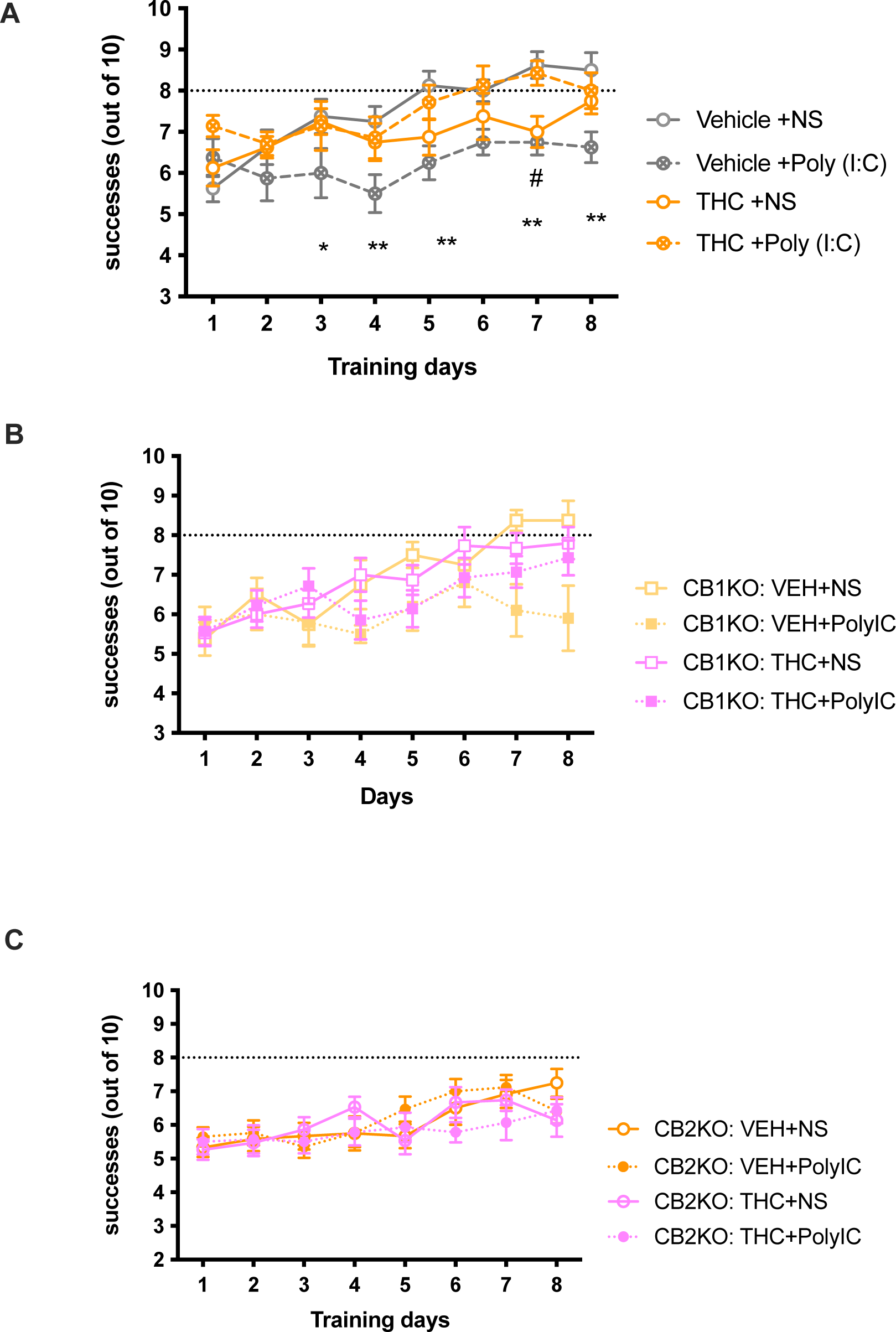
Maternal THC or MIA, but not their combination, slows learning in a working memory task in female offspring. **A.** Female offspring of wildtype dams treated with either THC or MIA learned the T maze task more slowly, but offspring of mice receiving THC and subjected to MIA learned the task as quickly as vehicle-treated mice. **B.** Maternal THC no longer impaired T maze learning in female CB1 KO offspring. MIA does not impair learning in CB1 KO offspring and combining THC and MIA rescued impairment. **C.** Vehicle treated CB2 knockout mice did not learn the working memory task to meet the standard criteria. Maternal THC and MIA impaired T maze performance and THC no longer rescued performance in CB2 knockout mice. T maze learning was assessed by 8-days of daily training with a fixed delayed and success rate recorded as a learning curve. Open symbols represent male offspring from dams receiving normal saline and solid symbols represent male offspring from dams receiving poly (I:C), both at GD16.5. Sample size in panel A: 8 for wildtype vehicle/ns, 8 for wildtype vehicle/poly(I:C), 8 for wildtype THC/ns, 9 for wildtype THC/poly(I:C) ;in panel B: 5 for wildtype vehicle/ns, 4 for wildtype vehicle/poly(I:C), 10 for CB1KO vehicle/ns, 8 for CB1KO vehicle/poly(I:C), 13 for CB1KO THC/ns, 14 for CB1KO THC/poly(I:C); in panel C: 12 for wildtype vehicle/ns, 5 for wildtype vehicle/poly(I:C), 13 for CB2KO vehicle/ns, 14 for CB2KO vehicle/poly(I:C), 9 for CB2KO THC/ns, 11 for CB2KO THC/poly(I:C)).*, p<0.05; **, p<0.01 compared with chronic vehicle and GD16.5 normal saline; #, p<0.05 compared with chronic vehicle and Poly (I:C) at GD16.5.

To test if CB1 receptors were required for MIA or PCE to impair working memory, we repeated the above treatments in CB1 KO mice. Here, we found that all CB1 KO male mice improved in task performance during training, irrespective of the perinatal treatment regimen (Figure 3B, F (5.725, 234.7) =18.4, P<0.0001), but there were significant differences between treatments (F (3, 41) =5.432, P=0.0031). MIA still impaired performance in CB1 KO mice (F [1, 16] =11.53, P=0.0037). However, CB1 receptors are required for PCE-induced deficits, as CB1 KO PCE and vehicle mice learned the task similarly (F [1, 21] =3.497, P=0.0755). As with wildtype mice, combining MIA+PCE treatment in CB1 KO mice did not affect learning of this task (F [1, 22] =3.650, P=0.0692). CB1 KO female mice improved their performance in the task during daily training (Figure 4B, F [5.857, 251.8], P<0.0001). Once again, there were significant differences between the groups (F [3, 43] = 3.539, P=0.0223). Specifically, female CB1 KO mice in the MIA group were significantly impaired compared with control mice (F [1, 16] = 5.193, P=0.0367). Impairment was not seen in female CB1 KO mice in the PCE group (F [1, 21] = 0.3432, P=0.5642). However, combined MIA + PCE was still protective in female CB1 KO mice (F [1, 20] = 2.938, P=0.1020). These results demonstrate that CB1 receptors are essential for PCE impairment in learning this task, but has, at most, a minor role in MIA or MIA + PCE effects.

In CB2KO mice, all four groups improved performance during training (Figure 3C, F (5.685, 244.5) = 7.544, P<0.0001), with no significance differences between treatment groups (F (3, 43) =1.679, P=0.1856), perhaps because of a slower rate of learning in the CB2 KO mice, limiting conclusions that can be drawn from these studies. In CB2KO female mice, although all the mice showed learning during daily training (Figure 4C, F [5.702, 307.9] = 6.729, P<0.0001), there was no difference between different treatment groups by 2-way ANOVA analysis (F [3, 54] = 0.9530, P=0.4217).

### 3.4. Effect of PCE and MIA on the delayed alternating memory test

After observing the impacts of PCE and MIA on working memory, we next examined treatment effect on the persistence of working memory. Thus, the day after completing the T maze memory learning phase, we ran another experiment where we progressively extended the delay between the forced and free choice trials in the alternation task to determine the impact of increasing delay on performance. For wildtype male mice, although performance decreased with increasing delay (Figure 5A, F (3.567, 103.4) =16.99, P<0.0001), there were significant differences between treatments (F (3, 29) =13.26, P<0.0001). Looking into each group’s difference, the MIA group showed the largest deficit with increasing delay (training, P=0.0016,95% C.I.= [0.9693, 3.114]; 10s, P=0.0169,95% C.I.= [0.4026, 4.097]), and became severe after prolonged delayed (15s, P=0.0007, 95% C.I.= [1.527, 4.973]; 30s, P=0.0025, 95% = [1.211, 5.286]; 60s, P<0.0001, 95% C.I.= [2.400, 5.600]). The PCE group was similarly impaired (training, P=0.0002,95% C.I.= [0.9693, 3.114]) and (10s, P= 0.0169, 95% C.I.= [0.4026, 4.097]; 15s, P=0.0420, 95% C.I.= [0.08139, 4.419]; 30s, P=0.0025, 95% C.I.= [1.211, 5.289]; 60s, P=0.0097, 95% C.I.= [0.9574, 6.043]). In perinatal combined treated mice, the success rate decreased at a rate similar to control mice. In wildtype female mice, performance decreased as delay increased (Figure 6A, F [4, 108] = 13.03, P<0.0001) and the rate of decrease depended on treatment (F [3, 27] = 10.51, P<0.0001). The MIA group showed impairment with extended delayed time (15s, P<0.0001,95% C.I.= [1.855, 6.145]; 30s, P<0.0001,95% C.I.= [2.105, 6.395]; 60s, P=0.0033, 95% C.I.= [0.8545, 5.145]). There were no significant differences between the other groups, though THC PCE groups trended towards impaired performance.

**Figure 5.**
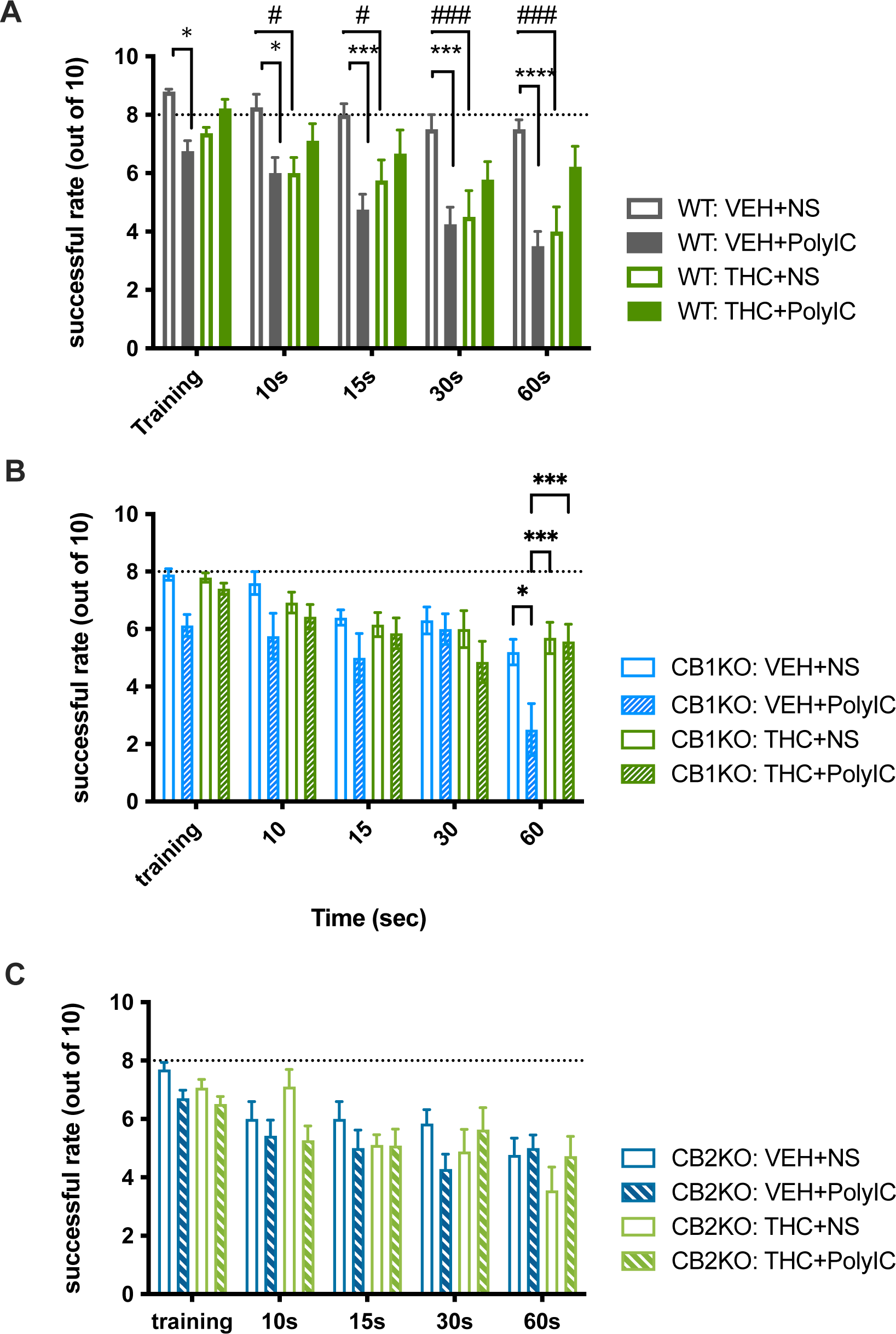
Maternal THC or MIA impairs memory consolidation, while the combination results in less impairment in male offspring. Memory consolidation was tested by increasing the delay time in the T maze task after 8 days of training. **A.** Maternal THC or MIA significantly decreased the number of successful choice as the delay time increased compared to both vehicle/saline and combined THC and MIA. While consolidation was impaired in both CB1 and CB2 KO mice, THC, MIA, or their combination had minor further effects in **(B.)** CB1 KO or **(C.)** CB2 KO mice. Open bars represent male offspring from dams receiving normal saline and bars represent male offspring from dams receiving poly (I:C), both at GD16.5. Sample size in panel A: 8 for wildtype vehicle/ns, 8 for wildtype vehicle/poly(I:C), 8 for wildtype THC/ns, 7 for wildtype THC/poly(I:C) ;in panel B: 3 for wildtype vehicle/ns, 4 for wildtype vehicle/poly(I:C), 8 for CB1KO vehicle/ns, 10 for CB1KO vehicle/poly(I:C), 15 for CB1KO THC/ns, 14 for CB1KO THC/poly(I:C); in panel C: 10 for wildtype vehicle/ns, 5 for wildtype vehicle/poly(I:C), 12 for CB2KO vehicle/ns, 17 for CB2KO vehicle/poly(I:C), 15 for CB2KO THC/ns, 14 for CB2KO THC/poly(I:C)).*, p<0.05; ***, p<0.001 compared with chronic vehicle and normal saline at GD16.5; #, p<0.05; ###, p<0.001 compared with chronic vehicle and Poly (I:C) at GD16.5.

**Figure 6.**
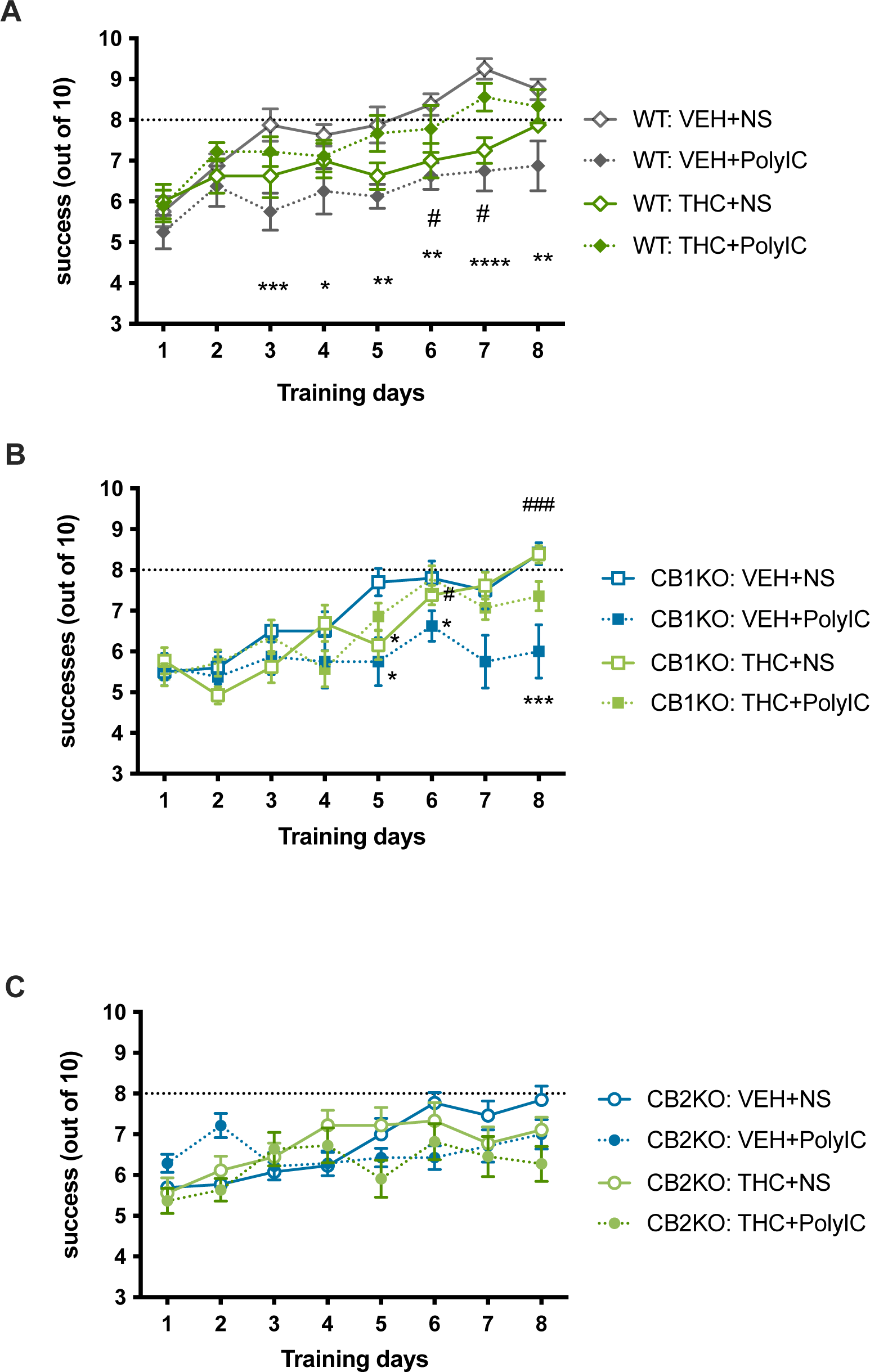
Maternal THC or MIA impairs memory consolidation, while the combination results in less impairment in female offspring. Memory consolidation was tested by increasing the delay time in the T maze task after 8 days of training. **A.** Maternal THC or MIA significantly decreased the number of successful choice as the delay time increased compared to both vehicle/saline and combined THC and MIA. While consolidation was impaired in both CB1 and CB2 KO mice, THC, MIA, or their combination had minor further effects in **(B.)** CB1 KO or **(C.)** CB2 KO mice. Open bars represent female offspring from dams receiving normal saline and bars represent female offspring from dams receiving poly (I:C), both at GD16.5. Sample size in panel A: 8 for wildtype vehicle/ns, 8 for wildtype vehicle/poly(I:C), 8 for wildtype THC/ns, 7 for wildtype THC/poly(I:C) ;in panel B: 3 for wildtype vehicle/ns, 4 for wildtype vehicle/poly(I:C), 8 for CB1KO vehicle/ns, 10 for CB1KO vehicle/poly(I:C), 15 for CB1KO THC/ns, 14 for CB1KO THC/poly(I:C); in panel C: 10 for wildtype vehicle/ns, 5 for wildtype vehicle/poly(I:C), 12 for CB2KO vehicle/ns, 17 for CB2KO vehicle/poly(I:C), 15 for CB2KO THC/ns, 14 for CB2KO THC/poly(I:C)).*, p<0.05; **, p<0.01; ****, p<0.0001 compared with chronic vehicle and normal saline at GD16.5.

Male CB1KO mice performance significantly decreased as delay increased (Figure 5B, F[3.257, 133.6]=16.12, P<0.0001), with significant differences between treatment groups (F[3, 41]=4.557, P=0.0076). But only a single difference emerged with multiple comparisons: CB1KO control and MIA group in training phase (P=0.0048, 95% C.I.= [-0.5819, 4.282]), which might occur if CB1 receptors are required to efficiently retain short term memory. Female CB1KO mice showed a strong decrease in performance with increasing delay (Figure 6B, F [3.547, 152.5] = 22.94, P<0.0001), with differences among treatment group (F[3, 43]=5.964, P=0.0017). Unexpectedly, the CB1KO female control group performed poorly with increasing delay, so there was no difference when compared with the MIA group, however the PCE group showed a deficit at 10s delay (P=0.0197, 95% C.I.= [0.2856, 3.548]) and the combined group was significantly impaired at the longest delay (60s, P=0.0133, 95% C.I.= [0.6433, 5.285]).

Male CB2KO mice performance significantly decreased with increasing delay (Figure 5C, F [3.393, 145.9] =13.57, P<0.0001), however there were no differences between treatments (F[3, 43]=1.415, P=0.2514). Similarly in female CB2KO mice, performance decreased with increasing delay (Figure 6C, F [3.477, 187.7] = 33.04, P<0.0001), with no significance differences among treatments (F [3, 54] = 1.373, P=0.2608).

## 4. Discussion

Environmental factors can adversely affect neurodevelopment, with serious consequences for the offspring. Since both MIA and PCE negatively affect neurodevelopment, we were interested in their combined impact. To examine this, we designed a mouse experimental paradigm, informed by human neurodevelopmental milestones (17,36), of a second trimester viral infection in a cannabis-using mother. To do this, we injected THC from GD5 until P10, which would correspond (in terms of cortical development) to consumption throughout pregnancy, and gave a single 5 mg/kg i.v. injection of poly (I:C) on GD16.5, mimicking severe viral infection.

Neither treatment nor genotype significantly affected activity or anxiety, as measured by the open field assay or elevated plus maze, respectively (Table 1). As expected, MIA or PCE given individually impaired novel object recognition and working memory (Figures 1-4), with more consistent deficits observed in males (Figures 1 and 3). Surprisingly, we found if we administered poly (I:C) at GD16.5 to dams receiving THC (PCE + MIA), NOR and establishment and consolidation of working memory were indistinguishable from control offspring in both sexes. Thus, the combination of PCE and MIA is protective against the deficits elicited by either treatment. These results are summarized in Figure 7.

**Figure 7.**
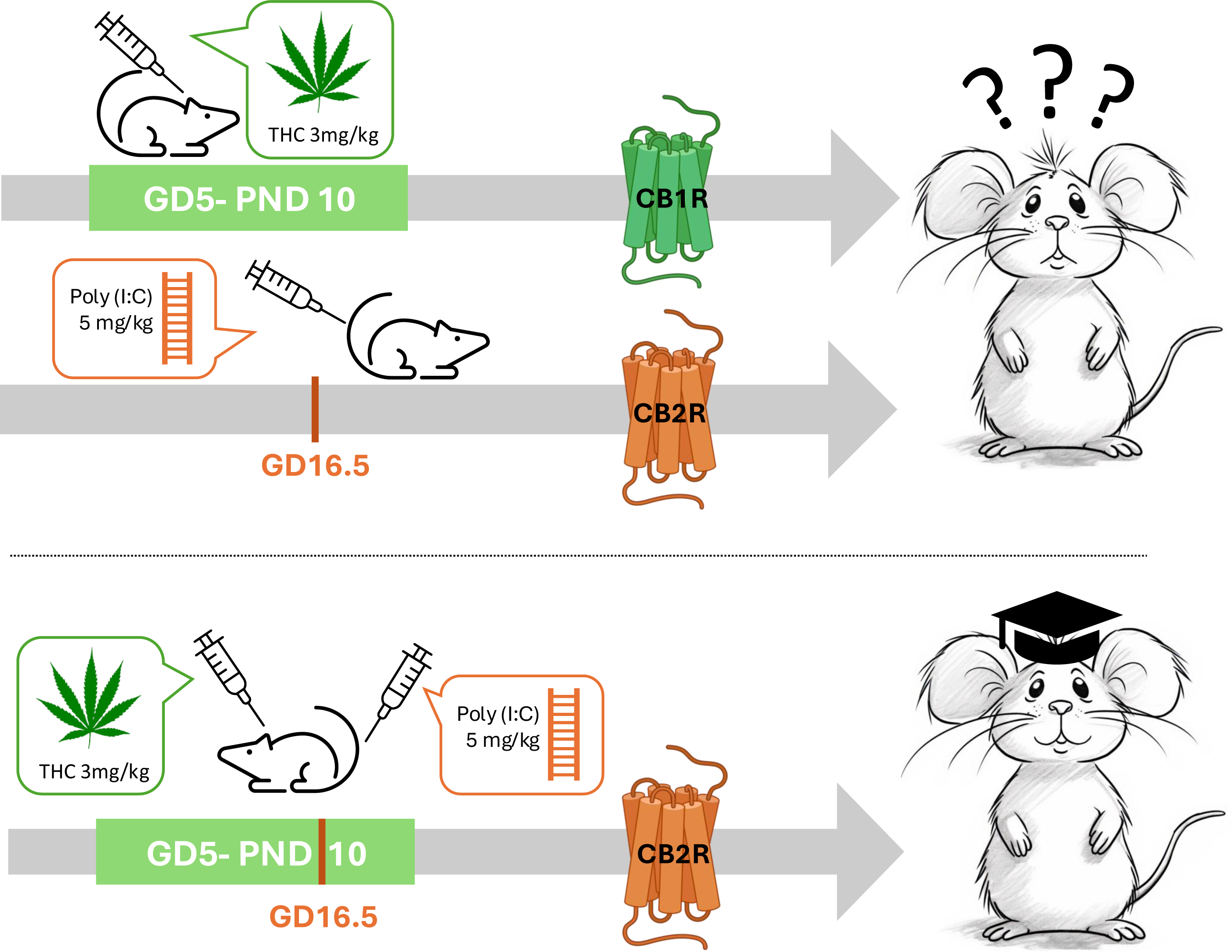
Graphical summary of the effects of PCE, MIA, and the combination. Maternal exposure to THC from gestational day 5 (GD5) to postnatal day 10 (PND10) results in cognitive dysfunction in offspring, mediated by CB1 receptors. Conversely, CB2 receptor activation is required for poly(I:C)-induced MIA-related cognitive deficits. Remarkably, when both perinatal exposures are combined, minimal cognitive impairment is observed in adult offspring, an effect mediated by CB2 receptors.

To identify which cannabinoid receptors were involved in the PCE deficits, we used transgenic mice lacking CB1 or CB2 receptors. We found that PCE essentially eliminated novel objection recognition in wildtype and CB2KO but not in CB1KO offspring, indicating that CB1 receptors play an important role in the recognition deficits elicited by PCE. In addition, PCE-treated CB1KO mice showed similar learning speed and no difference from wildtype mice as working memory was established, while PCE CB2KO mice still showed deficiencies in these measures. This result implicates CB1 receptors in cognitive deficits following PCE, while CB2 receptors don’t appear to have a role. In contrast, mice lacking CB2 receptors had normal novel object recognition and working memory consolidation after MIA (Table 2), suggesting that activation of CB2 receptors during MIA helps mediate MIA-induced deficits. Finally, the protection afforded by poly (I:C) on GD16.5 against the PCE-induced cognitive effects required CB2 receptors (Figure 1 and 3E-F).

**Table 2.** Summary of the impacts of MIA, PCE, and their combination on behaviors by genotype.

|  | wildtype |  |  | CB1KO |  |  | CB2KO |  |  |
| --- | --- | --- | --- | --- | --- | --- | --- | --- | --- |
|  | Novel object recognition | Working Memory: acquisition | Working Memory: consolidation | Novel object recognition | Working Memory: acquisition | Working Memory: consolidation | Novel object recognition | Working Memory: acquisition | Working Memory: consolidation |
| Control | ↑ | ↑ | ↑ | ↑ | ↑ | ↑ | ↑ | - | ↑ |
| MIA | ↓ | ↓ | ↓ | ↑/-<sup>a</sup> | ↓ | ↓ | ↑ | - | ↑ |
| PCE | ↓ | ↓ | ↓ | ↑ | ↑ | ↑ | ↓ | - | ↓ |
| PCE+MIA | ↑ | ↑ | ↑ | ↑/-<sup>a</sup> | - | - | ↓ | - | ↑ |

↑, increased discrimination performance/learning memory; ↓, decreased discrimination performance/learning memory; -, increased but either not significantly or did not achieve predefined standard (i.e., 8 successful trials for 3 consecutive days). a, female difference did not reach significance.

What might be the mechanism underlying this novel finding for a protective role of MIA in PCE? In the embryonic mouse brain TLR3, the target of poly(I:C) (37), is highly expressed in microglia and endothelial cells, and to a lesser extent on astrocytes and neurons(38). CB2 receptors are abundantly expressed on microglia and deletion of CB2 impairs TLR3 signaling (39). Thus, the combination of THC-engaged CB2 receptors and poly(I:C) activated TLR3 may protect against CB1-mediated PCE memory deficits by altering microglial responsiveness. Conversely, downregulation of microglial CB2 receptors by chronic THC during PCE may eliminate the detrimental effects of TLR3 activation in MIA. A microglial-mediated mechanism is also supported by the observation that PCE leads to neuroinflammation, signs of microglial activation, and impairment of endocannabinoid-mediated synaptic plasticity that extend at least into adolescence (40). It would be interesting to determine if mice subjected to PCE/MIA as in the current study demonstrate the same abnormal responses to THC as found in (40).

Embryonic endothelial cells also express substantial levels of TLR3 as well as CB2. Poly(I:C) - induced MIA increases blood brain barrier (BBB) permeability (41). In mature animals, CB2 activation reduces BBB permeability by decreasing NF-kB signaling (42), however it is not known if CB2 agonists have this effect in embryonic brain endothelial cells. Nonetheless, another potential explanation of the protective effects of THC is at the level of engagement of CB2 at the BBB.

Sex differences are now understood to be an important consideration in developmental studies. In the NOR task, male wildtype offspring receiving vehicle treatment reliably distinguish the novel from the familiar object. But the female vehicle group learned this task less well under the conditions of this study (e.g., Figures 2 and 4). If we combined all vehicle wildtype females tested to maximize statistical power, their recognition ability increased significantly after training (F (26) = 11.93, P=0.0011, 95% C.I. = [-0.4933, -0.1514]) and individual DI differences also showed an increased discrimination ability (mean= 0.2700, t [36] =5.125, P<0.0001, 95% C.I. = [0.1632, 0.376]), indicating wildtype female mice may learn this task less reliably than males under our experimental conditions. However, the differences in average performance to discriminate between the two objects was not statistically significant between sexes (t [61.77] =1.527, P=0.1319, 95% C.I. = [-0.2428, 0.0325]) limiting the conclusions that can be drawn. Overall, treatment effects were less pronounced in female mice, with the vehicle control group displaying more variability in performing most behaviors when compared to males. This variability introduces challenges in conclusively determining a treatment effect. However, the consistency observed within the same litter suggests that the estrous cycle may be a contributing factor. Future studies could incorporate estrous cycle monitoring (vaginal cytology) to control for this variable and improve the precision of the findings.

A potential confound in our experimental design comes from the ethanol in the vehicle. In preparing the injection suspension, the THC is first dissolved in ethanol. Mice received 10 μl of injectate/gm body weight. Thus, mice received, 0.5 μl ethanol/gm, or 0.39 mg ethanol/gm (0.39 g/kg). Studies of low dose experimenter administered ethanol typically use 2-3 g/kg, considerably higher than the mice will encounter here. Another way to look at this dose is through a low dose self-administration experiment in CD1 mice (43). In this experiment, blood alcohol concentrations (BACs) of 110-150 mg/dl were achieved and mild behavioral and anatomical abnormalities were observed. From the literature (44), a dose of 0.4 gm/kg of ethanol i.p. in male mice is expected to give a peak BAC of ∼20 mg/dl, with BAC returning to ∼0 within 60 minutes. Thus, while conclusively eliminating the possibility that the ethanol used here has a main effect will require the comparison of ethanol and non-ethanol vehicles, this possibility seems unlikely.

A related concern is that ethanol in the vehicle may be innocuous, but when combined with the low dose of THC, may produce deleterious synergism. Potential synergism between THC and ethanol effects on neurodevelopment has been examined in several studies (45). For example, combining inhaled THC (peak plasma levels ∼60 nM) with vaporized ethanol (peak plasma levels ∼175 mg/dl) to pregnant rat dams increased open field activity (OFA) in male offspring above levels seen with either drug alone (46). We did not observe an increase in activity in male mice receiving THC in the OFA in the current study (Table 1). This is consistent with the doses of ethanol and THC the mice received in the current, which would give projected plasma levels of 20 mg/dl and ∼6 nM for ethanol and THC (47), respectively, much lower than in the study cited above. Thus, while synergism between ethanol and THC is an important topic because the two drugs are frequently used together, it seems unlikely to be affecting our results.

### Concluding remarks and future directions

The interaction between THC and MIA provides indirect insight into when THC affects brain development relevant for the behaviors examined here. THC was given from GD5 to PND10. Since MIA given on GD16.5 was protective, this suggests that THC adversely effects brain development (relative to the behaviors measured in this study) at GD16.5 or later. This is consistent with human data that suggests cannabis use is most detrimental when continued throughout pregnancy (48), as GD16.5 roughly corresponds to second trimester human cortical development(49).

An interesting and clinically relevant question rising from our study is if MIA may protect from the adverse effects of human THC PCE. The motivation of this study was to determine if the individual adverse neurodevelopmental effects of MIA and THC PCE synergized. Much to our surprise, we found a protective effect. Poly(I:C) classically mimics infection by double stranded (ds) RNA viruses. There are few dsRNA viruses that are pathogens in humans, with rotaviruses being a notable exception. Future studies should determine if this effect is time specific, as MIA with poly(I:C) has different consequences depending on gestational age of poly(I:C) administration (50). In addition, it will be interesting to determine if other types of MIA, such as lipopolysaccharide (LPS), which is primarily thought to activate TLR4 (51), is similarly protective. Our study highlights a complicated and interesting interaction between MIA and PCE. Although individually, MIA or PCE is harmful for the offspring’s cognitive capability, their combination is protective. Whether a similar protective relationship is present in humans requires further investigation.

## Acknowledgements

THC was generously provided by NIDA Drug Supply. We would like to express our gratitude to Calan Quate for overseeing the breeding of the mouse colony, to Jim Wager-Miller for his technical support, and Brady Atwood for helpful conversations. We also extend our appreciation to Lab Animal Resource personnel for their diligent care of the mice and to the PBS Technical Support Staff for their assistance in designing and building some of the behavioral testing equipment.

## Author Contributions

H-TC and KM designed experiments, analyzed data, and wrote the paper. H-TC conducted experiments.

